# Engineered human lymph node stroma model for examining interstitial fluid flow and T cell egress

**DOI:** 10.1101/2024.12.03.622729

**Authors:** Jennifer H Hammel, Abhinav Arneja, Jessica Cunningham, Maosen Wang, Sophia Schumaecker, Yamilet Macias Orihuela, Tochukwu Ozulumba, Jonathan Zatorski, Thomas J Braciale, Chance John Luckey, Rebecca R Pompano, Jennifer M Munson

## Abstract

The lymph node (LN) performs essential roles in immunosurveillance throughout the body. Developing in vitro models of this key tissue is of great importance to enhancing physiological relevance in immunoengineering. The LN consists of stromal populations and immune cells, which are highly organized and bathed in constant interstitial flow. The stroma, notably the fibroblastic reticular cells (FRCs) and the lymphatic endothelial cells (LECs), play crucial roles in guiding T cell migration and are known to be sensitive to fluid flow. During inflammation, interstitial fluid flow rates drastically increase in the LN. It is unknown how these altered flow rates impact crosstalk and cell behavior in the LN, and most existing *in vitro* models focus on the interactions between T cells, B cells, and dendritic cells rather than with the stroma. To address this gap, we developed a human engineered model of the LN stroma consisting of FRC-laden hydrogel above a monolayer of LECs in a tissue culture insert with gravity-driven interstitial flow. We found that FRCs had enhanced coverage and proliferation in response to high flow rates, while LECs experienced decreased barrier integrity. We added CD4+ and CD8+ T cells and found that their egress was significantly decreased in the presence of interstitial flow, regardless of magnitude. Interestingly, 3.0 µm/s flow, but not 0.8 µm/s flow, correlated with enhanced inflammatory cytokine secretion in the LN stroma. Overall, we demonstrate that interstitial flow is an essential consideration in the lymph node for modulating LN stroma morphology, T cell migration, and inflammation.

## Introduction

Developing human immunocompetent engineered models is a crucial next step toward understanding disease progression and enhancing pre-clinical drug screening (1). Lymph nodes (LNs) are an essential part of the immune system, providing immunosurveillance of the lymph and mounting antigen-specific responses. Thus, immunocompetent models of the LN potentially have broad applicability in areas such as anti-tumor immunity, viral infections and vaccinations, and autoimmune disorders. Within the LN, a network of fibroblastic reticular cells (FRCs) guides T cell migration and enwraps conduits made of extracellular matrix to guide lymph fluid through the node for screening (2). Afferent and efferent lymphatic vessels, made up of lymphatic endothelial cells (LECs), regulate T cell entry and exit from the node (3). The LN further contains densely packed immune cells, including helper CD4+ T cells and cytotoxic CD8+ T cells. To provide immunosurveillance, T cells consistently recirculate from the blood to LN and back to blood on the scale of approximately 12 to 24 hours (4–6). Within the LN, T cell migration is guided by FRCs via direct contact and chemokine secretion. If a naïve T cell does not encounter an antigen, it leaves the LN through the efferent lymphatics and continues to circulate. This constant turnover allows for rapid detection of antigens. However, the contribution of physical forces, such as interstitial fluid flow, on the dynamics of T cell recirculation is not well understood.

The LN is organized to receive and sample lymph fluid, resulting in constant interstitial fluid flow (IFF) that bathes both the stroma and T cells. After filtering through the FRC-mediated conduit system, fluid subsequently leaves the lymph node through transmural flow to the efferent lymphatics. The rate of lymph flow drastically increases during inflammation (7–9), where the inflammatory state also reduces T cell egress. The impact of IFF on fibroblasts outside the lymph node is well established, with increased alignment (10–12), proliferation (13,14), motility (15,16), and upregulation of matrix related genes and matrix remodeling (14,17,18) all widely reported. However, only one study has observed LN FRC response to IFF, demonstrating enhanced networking and gene expression for chemokines (19). Additionally, lymphatic endothelial cell monolayers show enhanced permeability in response to transmural flow (20), mediated via the loosening of “button” junctions. Though the impact of IFF on lymphocytes is understudied, shear stress has been shown to induce migration in the opposite direction of flow (21) and altered proliferation in T cells (22). Thus, enhanced IFF could modify the behavior of the stroma and T cells to promote the adaptive immune response. Understanding how biophysical cues (e.g. flow, stiffness, matrix density) impact immune cell migration upon arrival to the LN is key toward developing immunocompetent models. Development of lymph node stroma models that could answer these questions is underway, with spatially organized murine models (23) and human models focused on LECs (24) or FRCs (19,25–29). However, no human model has been developed that incorporates both of these stromal components and interstitial fluid flow.

Herein, we sought to develop a spatially organized model of the LN stroma to understand the impact of IFF on the stroma and on retention or egress of T cells. Our model consists of FRC-laden hydrogel, representing the T cell zone, above a monolayer of LECs on the underside of a tissue culture insert, representing an efferent lymphatic vessel. By combining collagen and hyaluronic acid, this hydrogel model is capable of generating IFF driven by a gravitational pressure head. CD4+ and CD8+ T cells can migrate through the FRC region and egress across the LEC monolayer. We demonstrate the potential for this model to provide a vital tool toward understanding the impact of the stromal compartment and fluid flow on recirculating T cell dynamics.

## Methods

### Cell culture

Human lymph node lymphatic endothelial cells (LECs) (Sciencell) were cultured on fibronectin coated flasks in VascuLife® VEGF-Mv Endothelial Complete media, containing 5% fetal bovine serum and additional supplements as provided by the supplier (Lifeline Cell Technology). Human lymph node fibroblasts (FRCs) were cultured on poly-L-lysine coated flasks in fibroblast medium with 10% fetal bovine serum and 5% fibroblast growth supplement (Sciencell). All cells were maintained at 5% CO^2^ and 37°C.

T cells were isolated from leukocyte reduction system cones (STEMCELL Technologies) using a density gradient separation followed by immunomagnetic negative selection for target populations (STEMCELL Technologies). Donor age and sex were provided by the vendor. T cells were maintained in Immunocult T Cell Expansion Medium supplemented with 10 ng/mL interleukin-7 (R&D), interleukin-2 (Gibco), or with Immunocult Human CD3/CD28 T cell activator (STEMCELL Technologies).

### PhotoHA/collagen hydrogels

Methacrylated hyaluronic acid (PhotoHA) (Advanced Biomatrix) was reconstituted at 10 mg/mL in sterile 1X phosphate buffered saline (PBS) and stored at 4°C for up to one month. The photoinitiator, lithium phenyl-2,4,6-trimethylbenzoylphosphinate (LAP, Sigma Aldrich), was reconstituted at 17 mg/mL in sterile 1X PBS and stored at 4°C for up to two weeks. A 0.5% collagen solution was made on ice using high concentration collagen I from rat tail (Corning), 1N NaOH, water, and 10x PBS. LAP was added to PhotoHA at 0.02 times the volume of PhotoHA. Collagen solution, PhotoHA solution, and cell culture media was combined to give a final concentration of 0.4% PhotoHA and 0.2% Collagen. The hydrogel solution was first photocrosslinked at 50 mW/cm^2^ for 45 seconds with 385 nm light (Thorlabs Collimated LED 1700 mA controlled by Thorlabs DC2200 High-Power 1-Channel LED Driver) and then placed in a 37°C incubator for 30 minutes for thermal crosslinking of collagen.

### Pressure head driven interstitial flow

To create interstitial fluid flow, media was applied atop the hydrogel to generate a pressure head as previously described (30,31). For static conditions, 700 μL of media was placed in the lower chamber of a 12 well plate, and 100 μL was placed atop the gel. For 0.8 μm/s flow, 100 μL of media was placed in the lower chamber of a 12 well plate, and 700 μL was placed atop the gel. For the high flow condition, 12 mm fluorinated ethylene propylene (FEP) clear tubing was cut and secured onto the tissue culture insert and 2.7 mL is placed atop the hydrogel. To accommodate tubing and maintain sterility, lid raisers in the dimensions of a 12-well plate were 3D printed and sterilized with ethanol.

### Flow-mediated egress assays

CD4+ and CD8+ T cells were labeled with 1 μM CellTrace™ CFSE (Invitrogen) and embedded in PhotoHA/collagen gel precursor prior to crosslinking as described above at 500,000 cells per mL of hydrogel. AIM-V universal media was used to create pressure heads, supplemented with IL-7 or IL-2 for naïve T cells and CD3/CD28 immunocult activator for activated T cells. After 24 hours, flow through media was collected and analyzed via flow cytometry.

### LN stroma model

At confluence, LECs were passaged with Accutase and seeded in Vasculife complete on the bottom of 12-mm tissue culture inserts with 8 µm pore size (Corning) at 50,000 per insert for 2 hours (32–34). Vasculife was added and the LECs were cultured for 48 hours to reach confluency. Next, FRCs were resuspended in PhotoHA-collagen solution formulated as described above and photo- and thermally crosslinked in the LEC-coated tissue culture inserts. The LN stroma model was cultured for 24 hours prior to experimental manipulation. For flow studies, pressure heads were applied as described above and LN models were cultured for 24 hours prior to fixation with 4% paraformaldehyde. For T cell studies, CD4+ and CD8+ T cells were labeled with 1 µm CFSE and 50,000 were added to each LN model in the media above the hydrogel under designated flow conditions. After 24 hours, flow-through media was collected for flow cytometry and models fixed for staining and imaging.

### Flow cytometry

Flow-through media was spun down at 0.3 rcf for 10 minutes. Then, cell pellets were resuspended in PBS and transferred to 96 well plates. For viability staining, LIVE/DEAD™ Fixable near IR reactive dye (Invitrogen) was diluted 1:1000. Cells were spun down at 2000 rcf for 1 minute and resuspended with viability dye for 15 minutes on ice. Cells were spun down again and washed with flow cytometry buffer consisting of Hank’s balanced salt solution (HBSS) + 3% bovine serum albumin (BSA) twice before flow cytometry was performed on a Guava Cytometer. Analysis is performed via GuavaSoft. Singlets were gated using forward scatter height and area. T cells were identified via viability and CFSE staining. The concentration of T cells was then extrapolated to an estimated total count to report percent egress.

### Proliferation assay

After 18 hours, 10 μM EdU was added to culture media for the final 6 hours. Gels were fixed in 4% formaldehyde in 1x PBS for 1 hour. Detection was performed using Click-iT™ EdU Alexa Fluor™ 488 dye as specified by the manufacturer (Invitrogen). The number of EdU+ nuclei and the total number of nuclei, as detected by DAPI, were hand quantified in FIJI to report % proliferation.

### Cytokine analysis

Media was flash frozen in liquid nitrogen and stored at −80°C until assays were performed. Hydrogels were removed from the tissue culture insert with forceps and degraded with collagenase and dispase for 30 minutes at 37°C while shaking. Degraded hydrogel solution was then stored at −80°C until use. Interferon gamma was detected using an enzyme-linked immunosorbent assay as specified by the manufacturer (BioLegend). A four parameter logistic curve was used to determine IFNgamma concentration in samples (myassays.com). IL-6, CCL2, CXCL8, and CXCL12 were detected using a ProCartaPlex Multiplex Immunoassay (ThermoFisher). Quantification was performed on a Luminex xMAP. A five parameter logistic curve was generated via ThermoFisher ProcartaPlex Analysis App to determine concentrations.

### Immunofluorescence

To visualize cell organization, we performed immunocytochemistry on fixed LN stroma models for F-actin and CD31. LN stroma models were fixed in 4% formaldehyde for 1 hour. Blocking with donkey serum in 0.01% Triton x-100 was performed for 6 hours at room temperature (RT) on a rocker. LN stroma models were then incubated with 4 µg/mL sheep anti-human CD31 (R&D) in blocking solution overnight at 4°C on a rocker. LN stroma models were rinsed with PBS three times for 20 minutes on a rocker at RT. Next, gels were incubated with 5 µg/mL donkey anti-sheep AlexaFluor 647 in blocking solution overnight at 4°C on a rocker, followed by a rinse as previously described. Finally, gels were stained with 1:400 Alexa Fluor™ 488 Phalloidin and DAPI for 6 hours at RT on a rocker. LN stroma models were rinsed and stored in PBS until imaging.

### Imaging

Immunostained LN stroma models were placed on a glass coverslip to allow imaging of the intact model. For whole-gel imaging, Z-stacks with a 10 µm step size were taken on a Nikon Spinning Disk with a 10X objective. For LEC monolayers, a 20X objective was used. Three regions of interest were captured for each technical replicate. For second harmonic generation, an LSM 880 confocal microscope and 20X objective were used as previously described (35).

### Image quantification

Image analysis was performed in FIJI (36). Total F-actin+ area was used to determine the FRC distribution throughout the model. CD31+ area was used to determine LEC coverage of the tissue culture insert. CD31+ junctions were analyzed for disruption by counting LECs with discontinuous cell-cell contact. Portions of the monolayer where LECs were missing due to death were not considered to be disrupted junctions. For T cell counts, FIJI’s analyze particle function was used.

### Gene expression

LN stroma constructs were removed from tissue culture inserts with forceps and placed into Trizol. RNA was isolated via phase separation and purified via Quick-RNA MicroPrep Kit including the optional DNase treatment (Zymo Research). Fifty ng of RNA was converted to cDNA using the QuantiTect Reverse Transcription Kit (Qiagen). Gene-specific TaqMan assays (ThermoFisher Scientific) were used to quantify expression (**Table S1**) using 2 µL of cDNA template and TaqMan Universal Master Mix (ThermoFisher Scientific) and cycled following the manufacturer’s instructions. PCR reactions were run in triplicate on a QuantStudio 3 (ThermoFisher Scientific). No template controls were included in each PCR run.

A standard curve of pooled samples was run for each gene to allow quantitative analysis. The quantity of each gene was determined from the standard curve, and normalized with the housekeeping gene, GAPDH, to provide relative expression for each sample.

### Tissue staining

Formalin-fixed paraffin embedded (FFPE) normal human lymph nodes (AMSBIO) were sectioned at 5 μm and adhered to glass slides. Multi-plex staining was performed according to the manufacturer’s specifications (Akoya Biosciences). Slides were imaged on an Akoya Phenocycler.

### Statistics

One-way or two-way analysis of variances (ANOVA), followed by Tukey’s multiple comparison t-tests were performed using GraphPad Prism version 10.0.0 for Windows (GraphPad Software, Boston, Massachusetts USA) with significance at p<0.05. All p-values are reported on the graphs.

## Results

### Lymph node stroma model development

In the lymph node, organized fibroblastic reticular cells orchestrate T cell migration. T cells circulate through the lymphatic system and travel through LNs, where FRCs siphon lymph and dendritic cells present antigens to naïve T cells (2). Once an antigen is detected, T cells become activated and the entire LN mounts an immune response, with antigen-specific immune cells egressing from efferent lymphatics (3). However, if that antigen is not found, the T cells will remain naïve and eventually egress through the same pathway (**Figure 1A**).

**Figure 1.**
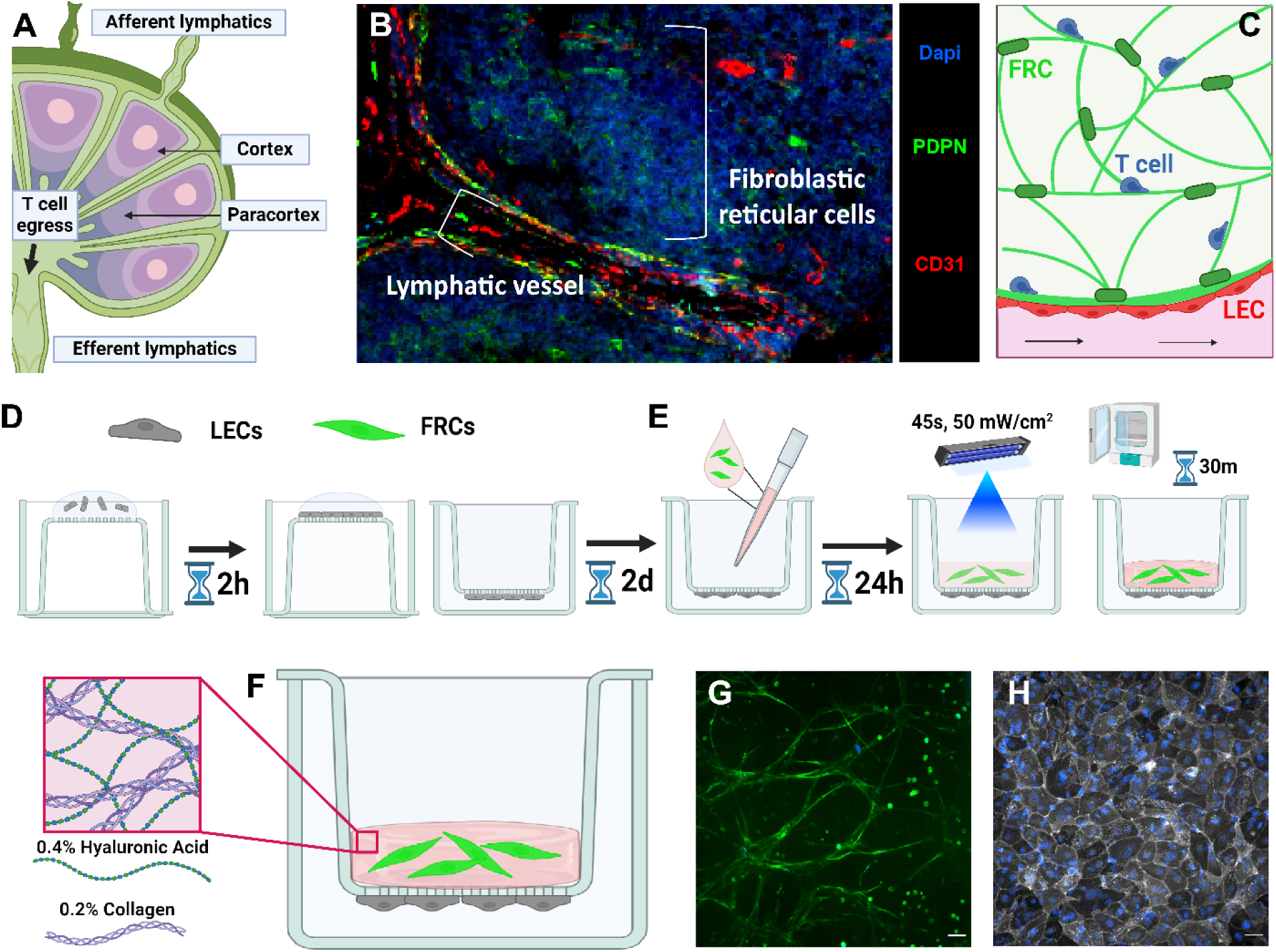
Lymph node stroma model development. We sought to develop a model of immune cell egress from the lymph node (A). In human lymph nodes, lymphatics (PDPN+,CD31+) and fibroblastic reticular cells (PDPN+) form the structure of the T cell zone (B). Schematic of intended structure of T cell zone model (C). LECs were seeded on the underside of a tissue culture insert (D). PhotoHA-collagen gels laden with FRCs are crosslinked above the LECs and then incubated for 30 minutes (E). After thermal crosslinking, the final gel consists of hyaluronic acid and collagen (F). With this methodology, FRCs formed networks (G) and LECs formed an intact monolayer (H). Scale bars are 50 microns.

We sought to develop a model to recapitulate the spatial organization of an FRC network and an efferent lymphatic vessel seen in human samples (**Figure 1B,C**). We utilized commercially available primary human FRCs and LECs derived from lymph nodes to enhance physiological relevance and adaptability. Using a tissue culture insert, intact LEC monolayers were formed on the underside (**Figure 1D**), oriented so that their basal side was in contact with the tissue culture insert and thus the FRC-laden hydrogel, while the apical side opened into the well. This orientation allowed the monolayer to serve as an efferent lymphatic. Within the insert, a FRC-laden hydrogel was photo- and thermally crosslinked above the LEC monolayer (**Figure 1E**), creating a spatially distinct LN stroma model (**Figure 1F**). Within, the FRCs formed inter-connected networks in the model (**Figure 1G**) and LECs maintained a confluent monolayer (**Figure 1H**). Thus, we recapitulated parts of the architecture of the T cell zone in a facile and simplified way.

This methodology was further compatible with magnetic resonance imaging (MRI), enabling a detailed analysis of the fluid flow through the model. We utilized dynamic contrast enhanced MRI to calculate the local (pixel-by-pixel) diffusion coefficient and the magnitude and direction of convective flow within the gel. The magnitude and direction result in a vector flow field, which is used to analyze the physical paths of fluid flow. These paths, visualized as streamlines, reveal the continuous trajectories that fluid particles follow through the gel matrix over time. In an empty PhotoHA-collagen gel, streamlines showed that flow is well distributed throughout the hydrogel (**Supplemental Figure 1D**). When LN stroma was present, the streamlines gathered and converged, with fewer locations of flow exiting the hydrogel (**Supplemental Figure 1E**). This is captured quantitatively in the significantly decreased divergence of velocity vectors with the LN stoma present **(Supplemental Figure 1F)**. This finding suggests that, even though the diffusion (**Supplemental Figure 1G**) and flow velocity (**Supplemental Figure 1H**) do not significantly change in the presence of LN stroma, the LN stroma reshapes the flow trajectories, indicating restricted or channeled movement of fluid through the matrix. Thus, we concluded that the LN stroma does alter fluid flow through the hydrogel.

### The LN stroma model predicts opposing responses of LECs and FRCs to interstitial fluid flow

We first utilized this model to characterize stromal cell response to IFF in the absence of lymphocytes. As we saw that the presence of LN stroma altered transport, we wished to examine morphological structures that formed within the model that could change paths of flow. Secondarily, we sought to determine how interstitial velocities could alter these cellular components. Thus, the LN stroma model was subjected to 0, 0.8, or 3.0 µm/s flow to probe IFF response (**Figure 2A**). These flow rates were accomplished via pressure head and validated via volumetric measurements (**Supplemental Figure 2**). To accommodate the 3.0 µm/s flow rate, tubing was secured onto the tissue culture insert (**Figure 2B**) and 3D printed lid raisers allowed for sterility (**Figure 2C**).

**Figure 2.**
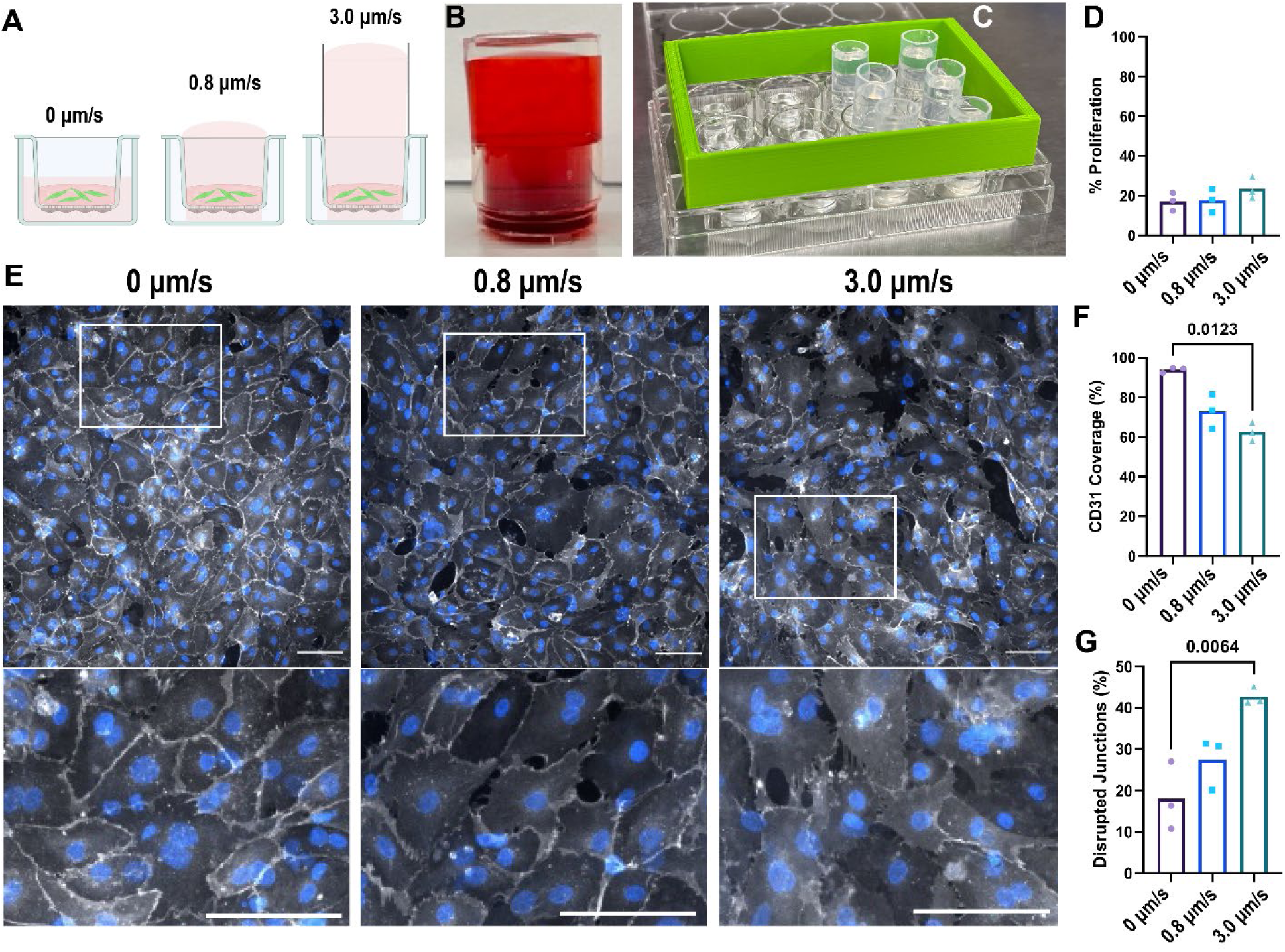
LEC barrier integrity is decreased under interstitial flow. Pressure heads of media are utilized to drive interstitial flow (A). To create the highest magnitude flow rate, PE50 tubing was secured onto the tissue culture insert with no leaks (B). 3D printed lid raisers to the specifications of a 12 well plate (C) maintain sterility in the incubator. LEC proliferation is unchanged by interstitial flow, as quantified by % of EdU+ cells (D). Representative images show LECs visualized with CD31 (gray). Nuclei are stained with DAPI (blue). Scale bars are 100 µm (E). LEC coverage (F) and disrupted junctions (G) are quantified. Each point represents a biological replicate from an independent experiment, for n=3. Each data point represents a biological replicate (n=3). Significance was determined by two-way ANOVA followed by Tukey’s t-test, with significant p values (<0.05) reported on each graph.

We first examined the LEC response to IFF. LEC proliferation was unchanged by IFF (**Figure 2D**). LEC monolayers demonstrated intact monolayers under static conditions and disruption under flow. Visually, LEC-LEC junctions showed loose, spindly connections, with small gaps in between under flow conditions (**Figure 2E**). Quantification demonstrated that LEC coverage was significantly decreased under 3.0 µm/s flow (**Figure 2F**). In addition, LEC junctions were significantly disrupted by 3.0 µm/s flow (**Figure 2G**).

Next, we sought to examine changes to FRCs within the bulk hydrogel in response to IFF. Representative images demonstrated FRC networks visualized with a phalloidin stain for F-actin (**Figure 3A**). FRC proliferation was significantly increased under 3.0 µm/s flow (**Figure 3B**). Similarly, FRC network area coverage was only significantly increased by 3.0 µm/s flow (**Figure 3C**). Next, we tested whether FRCs altered chemokine expression in response to IFF. We selected chemokines that are known to be involved in T cell migration and secreted by the primary fibroblastic reticular cells in the model (25) and examined expression at the gene level. While CXCL8 and TGFbeta had upward trends (**Supplemental Figure 3D,E**), only CCL2 stood out as significantly upregulated by 0.8 μm/s flow, but not 3.0 μm/s flow (**Figure 3D**).

**Figure 3.**
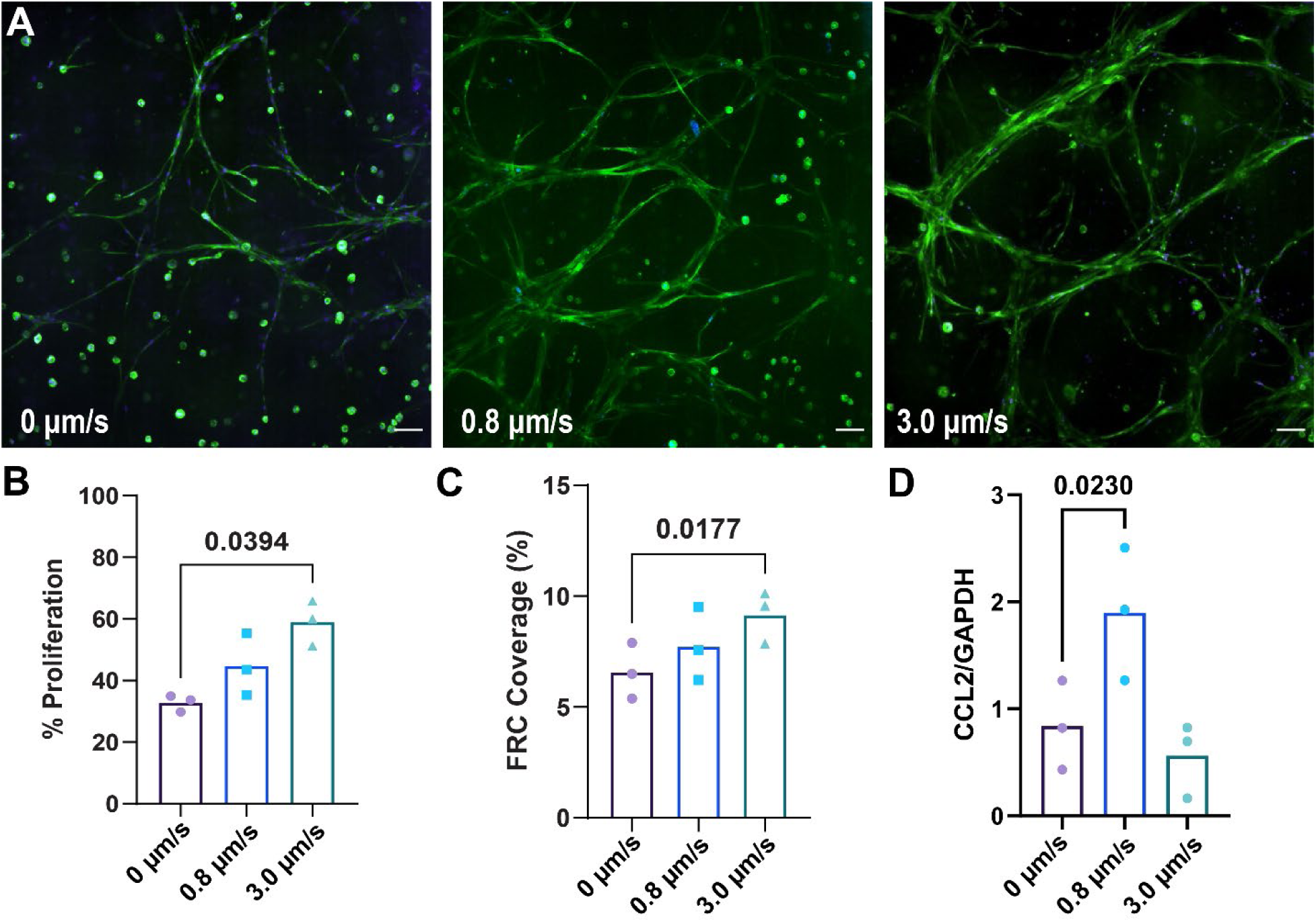
Fibroblastic reticular cells have enhanced proliferation and coverage in response to IFF. Representative images of FRCs under 0, 0.8, and 3.0 µm/s flow rates, visualized in green with F-actin (A). Scale bars are 100 µm. FRC proliferation (B) and coverage (C) are quantified. Gene expression of CCL2 is reported (D). Each data point represents a biological replicate (n=3). Significance was determined by one-way ANOVA followed by Tukey’s t-test, with significant p values (<0.05) reported on each graph.

To further understand changes occurring in the microenvironment, we examined the gene expression of matrix elements by FRCs in the LN stroma model under flow. Dense matrix is known to decrease T cell infiltration and motility (37–39), and could impact behavior in the lymph node. FRCs are known to secrete major structural collagens, such as type I collagen, basement membrane proteins, such as fibronectin, and proteolytic enzymes, such as matrix metalloproteinases (40,41), but the response to flow is unknown. We observed that the matrix remodeling related genes COL1A1 (**Supplemental Figure 3A**), FN1 (**Supplemental Figure 3B**), and MMP2 (**Supplemental Figure 3C**) were not significantly altered under 0.8 or 3.0 μm/s flow in the 24 hours of flow conditioning. Additionally, second harmonic generation imaging demonstrated no distinct visual differences in collagen structure between static and flow conditions (**Supplemental Figure 4**). As the alignment (42,43) and type of matrix deposited impacts T cell migration (44,45), future staining of matrix elements may reveal phenotypic changes outside of gene expression. Additionally, extending the time of culture may be necessary to sufficiently observe deposition.

Overall, these results demonstrated that IFF was disruptive to LEC coverage and junctions in the model, consistent with the known loosening of button junctions by efferent lymphatics to collect interstitial fluid (46). Furthermore, IFF was beneficial to FRC proliferation and coverage, particularly at higher flow rates, tracking with prior studies (19). Thus, this model successfully predicted the response to IFF for both cell types, and is ready for further exploration of these effects in the future, including for the effect of flow on matrix deposition and chemokine secretion.

### Activated T cell egress is decreased with interstitial fluid flow in monoculture

Interstitial fluid flow is an ever-present force within the body, yet the study of its influence on cell behavior in the immune system is still nascent. Next, we sought to understand the impact of interstitial fluid flow on T cell behavior in the LN microenvironment, where CD4+ and CD8+ T cells (**Supplemental Figure 5A**) experience drastic changes in interstitial fluid flow based on the level of inflammation. First, we tested how T cells alone respond to IFF. Naïve or activated T cells were encapsulated in PhotoHA/collagen gels and subjected to 0, 0.8 (physiological) or 3.0 µm/s (pathological) flow for 24 hours (**Supplemental Figure 5B).** Overall, we saw significantly less egress in naïve cells compared to their activated counterparts (**Supplemental Figure 5C**). Egress trended downward with interstitial fluid flow in activated CD4+ T cells and was significantly decreased in activated CD8+ T cells. It is important to note that photocrosslinking produces reactive oxygen species (ROS) that interfere with T cell migration (28), and thus, naïve T cells may be more susceptible to ROS than activated T cells.

Beyond migration, we also examined the secretion of IFNγ in response to flow. Interestingly, IFNγ secretion by activated CD8+ T cells was significantly increased by interstitial flow, both between the static control and between flow rates, demonstrating the potential importance of flow magnitude in this effect (**Supplemental Figure 5D**). Notably, IFNγ can bind to hyaluronic acid (47), and thus, exploration of IFNγ captured within the hydrogel could be of interest.

Flow-mediated changes to T cell egress in mono-culture could rely on a variety of factors. In the lymph node, T cells exhibit random motion as they scan for an antigen, and alternate between slow and fast migration (48). Naïve T cells and activated T cells have similar speeds in vivo, but activated T cells undergo more directed and environmentally guided motion, at times forming ‘swarms’ of stationary cells (48–50). In in vitro systems, T cells have been reported to migrate faster when given activating cues such as intracellular adhesion molecule-1 (51). Generally, changes to cell migration under flow in monoculture are often attributed to self-propelled chemotaxis via autologous gradients (52), or cells moving toward nutrients. Within monocultures, the decrease in T cell egress could potentially be caused by T cells moving against the direction of flow, as noted with shear stress (21), or moving toward the nutrients within the pressure head of media.

### T cells have reduced egress from the LN stroma under IFF

In vivo, T cell migration is mostly regulated by other cell types in the microenvironment. Therefore, we sought to examine T cell egress in the context of the full LN stroma model. After establishing the LN stroma model and culturing for 24 hours (**Figure 4A**), naïve or activated CD4+ and CD8+ T cells were added above the FRC-laden hydrogel under static or flow conditions to simulate T cell recirculation (**Figure 4B**). After 24 hours, flow-through media was collected and T cells were counted (**Figure 4C**). All T cell populations were able to migrate through and egress from the LN stroma model. Naïve and activated CD4+ T cells demonstrated similar % egress from the LN stroma model (**Figure 4D,E**). For both naïve and activated CD4+ T cells, egress was significantly decreased between static and flow conditions, but not between flow magnitudes. CD8+ T cells also exhibited similar levels of egress between naïve and activated (**Figure 4F,G**). For naïve CD8+ T cells, there were no significant differences in egress under flow. For activated CD8+ T cells, egress was decreased between static and flow conditions, but not between flow magnitudes.

**Figure 4.**
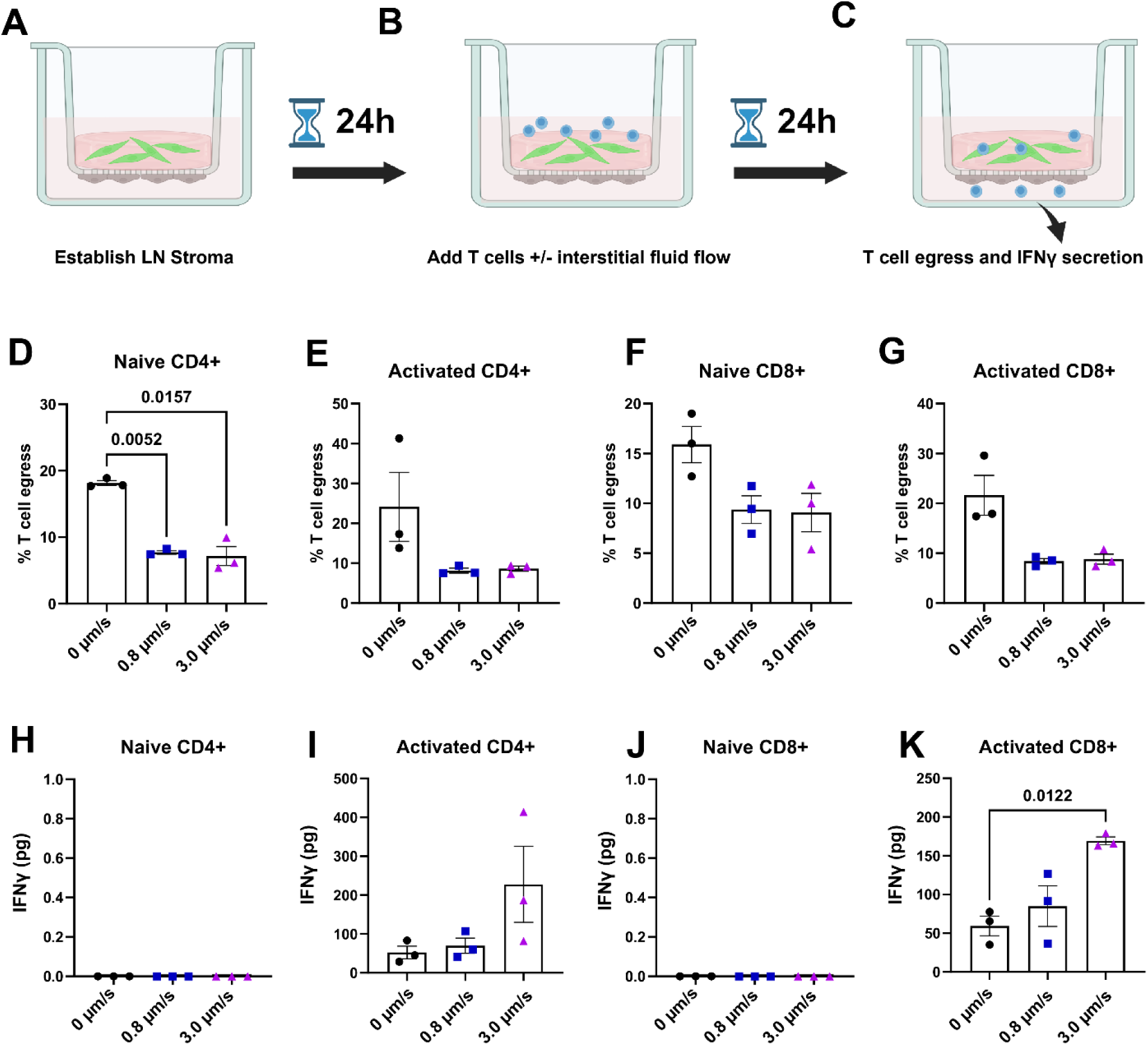
T cell egress through the LEC monolayer is decreased with interstitial fluid flow in the LN stroma. LN stroma models are established as previously described (A). After 24 hours, T cells with and without interstitial flow are added (B). After an additional 24 hours, media from underneath the model is collected and analyzed (C). Naïve CD4+ T cells have significantly decreased egress under both flow conditions (D). Activated CD4+ T cells (E), naïve CD8+ T cells (F), and activated CD8+ T cells (G), show that egress trends downward with flow. Interferon-gamma secretion was not detected by naïve CD4+ T cells (H). Activated CD4+ T cells secreted interferon-gamma that trended upward with 3.0 µm/s flow (I). Naïve CD8+ T cells did not secrete detectable levels of IFNγ (J). Activated CD8+ T cells secreted significantly more IFNγ under 3.0 µm/s flow compared to static conditions (K). Each data point represents a biological replicate and different T cell donor (n=3). Significance was determined by one-way ANOVA followed by Tukey’s t-test, with significant p values (<0.05) reported on each graph.

As noted, T cells produce IFNγ as a function of activation, and this cytokine has known impacts on the stroma. IFNγ is anti-lymphangiogenic (53,54), and FRCs respond to IFNγ by suppressing T cell proliferation (55,56). We quantified IFNγ in the supernatant to determine how IFF altered T cell cytokine secretion in the LN stroma and if that correlated with changes to the stroma. In the presence of LN stroma, naïve CD4+ T cells did not produce detectable levels of IFNγ (**Figure 4H**), whereas activated CD4+ T cells trended toward higher IFNγ secretion under flow (**Figure 4I**). Similarly, naïve CD8+ T cells did not produce detectable levels of IFNγ (**Figure 4J**), while activated CD8+ T cells secreted significantly more IFNγ under 3.0 μm/s flow (**Figure 4K**).

These results demonstrate that higher flow rates are required to alter IFNγ secretion in the presence of LN stroma. However, as naïve T cells do not secrete IFNγ, other cytokines are likely involved in altered egress. In the LN stroma model, the changes in T cell egress could still be due to nutrients. However, interstitial fluid flow increases with inflammation, which could signal a need for T cell response. Thus, this physical signal of flow could encourage T cells to be retained in the LN stroma model to search for antigen. Overall, these results demonstrate that T cells are responsive to IFF in the LN stroma.

### LEC barrier integrity is compromised in presence of T cells and flow

As the LN is a holistic system with continuous communications between cell types, we sought to examine how the presence of T cells and IFF would in turn impact the LN stroma. Representative images of the LEC monolayers demonstrated disruption in the presence of naïve and activated CD4+ and CD8+ T cells, which was exacerbated by IFF (**Figure 5A, Supplemental Figure 6**). The presence of IFF significantly decreased %CD31 coverage in the absence of T cells (**Figure 5B**). Interestingly, the presence of naïve CD4+ T cells seemed to mitigate LEC disruption, whereas with activated CD4+ T cells and naïve and activated CD8+ T cells, flow-mediated disruption was still present (**Figure 5B**). LEC junction disruption was significantly increased by flow conditions in the control, but not in the presence of naïve or activated CD4+ T cells or activated CD8+ T cells (**Figure 5C**). Overall, the presence of T cells erased the impacts of IFF in LEC junction disruption, but not overall coverage. Many T cell-secreted cytokines, including IFNγ, have anti-lymphangiogenic effects (53,54). Additionally, T cells are likely egressing between the junctions of LECs, increasing their disruption outside of flow magnitude.

**Figure 5.**
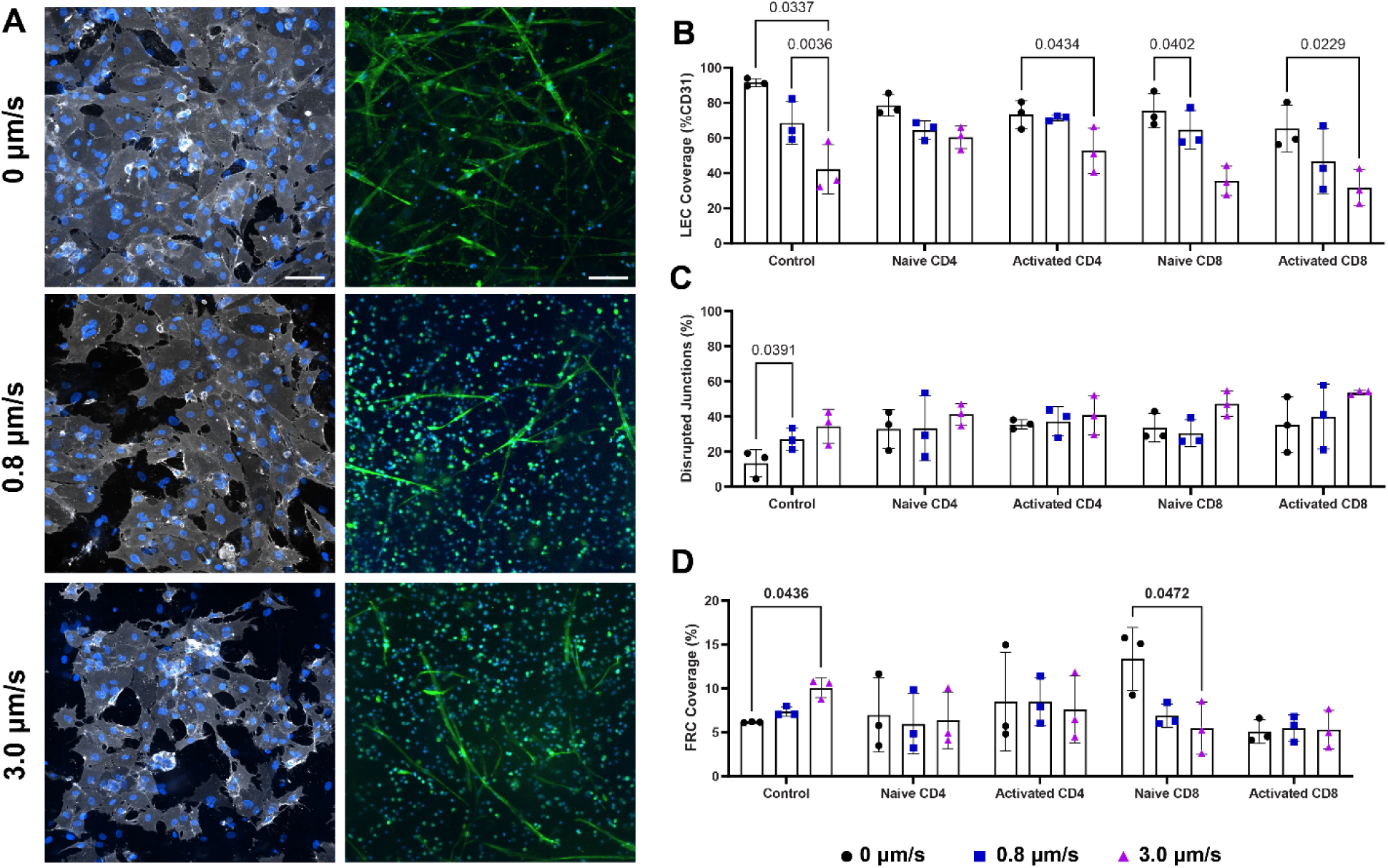
Presence of T cells disrupts LEC junctions regardless of flow and ameliorates flow-induced changes to FRCs. LECs are visualized in representative images with CD31 in gray and DAPI in blue (scale bar 100 microns), and FRCs are visualized with F-actin in green (scale bar 50 microns), in the presence of naïve CD8+ T cells (blue) (A). Quantification of LEC monolayer coverage (B) and disrupted junctions (C). FRC coverage in the presence of T cells and interstitial flow (D). Scale bar is 50 µm. Each data point represents a biological replicate (n=3). Significance was determined by two-way ANOVA followed by Tukey’s t-test, with significant p values (<0.05) reported on each graph.

### FRCs invasion into the LEC monolayer is decreased with T cell co-culture

As fibroblasts are known to be more migratory under flow (10,13–18), we sought to determine whether disruption of the LEC monolayer could be due to FRC invasion and whether T cells mitigated this effect. To do this, F-actin staining was quantified and compared to CD31 staining. Representative images show fibroblastic morphology of F-actin stains on the LEC monolayer, demonstrating that there is some level of FRC invasion into the LEC monolayer (**Supplemental Figure 7**). The percent area of F-actin was not significantly altered by interstitial flow in the control or CD4+ T cell groups (**Supplemental Figure 8A**). Interestingly, F-actin area was significantly decreased by flow in the presence of CD8+ T cells (**Supplemental Figure 8A**). To determine how much of the F-actin staining was coming from FRCs vs LECs, the ratio of F-actin to CD31 was quantified. A higher ratio indicates a higher number of FRCs compared to LECs (more invasion), as FRCs are only F-actin+ and LECs are double positive. In the absence of T cells, the ratio of F-actin to CD31 was significantly increased under 3.0 μm/s flow. Amongst all CD4+ and CD8+ T cells, the ratio was unchanged by flow (**Supplemental Figure 8B**), suggesting that FRCs are not displacing LECs as an effect of T cell co-culture. Overall, these results indicate that T cell presence mitigated flow-induced FRC invasion.

### Flow response of FRCs is mitigated by T cell presence

Next, we sought to examine how T cell presence would impact the FRC network. FRC coverage was significantly increased under 3.0 μm/s flow in the absence of T cells (**Figure 5D**). Interestingly, this effect of flow was prevented by the presence of naïve and activated CD4+ T cells as well as activated CD8+ T cells, and was reversed by the presence of naïve CD8+ T cells. Representative images show CD4+ and CD8+ T cells with FRCs in the model (**Figure 5A, Supplemental Figure 9**). Thus, these results demonstrate that just as FRCs support T cell function, the T cells in turn may be secreting factors that alter LN stroma response to flow. For example, lymphotoxin secretion by T cells has been shown to support FRCs (57). In addition, a hallmark of activated T cells is the secretion of IFNγ that initiates a feedback loop with FRCs meant to maintain LN function (55). These results show that T cells mitigate or reverse the flow response of FRCs, suggesting an important crosstalk pathway for future exploration.

### Inflammatory cytokine secretion is enhanced by 3.0 µm/s flow

Finally, to understand why T cell egress and stroma morphology were altered, we sought to determine whether the presence of IFF was inducing an inflammatory state in the LN stroma. We quantified inflammatory chemokine secretion under 0, 0.8, and 3 µm/s flow in LN stroma with and without CD4+ T cells.

Overall, there were no significant differences between LN stroma alone and LN stroma with CD4+ T cells. Above, we reported that CCL2 gene expression was upregulated by 0.8 µm/s flow. Interestingly, this was not seen at the protein level. CCL2 secretion in the LN stroma was upregulated under 3.0 µm/s compared to 0.8 µm/s flow, but not the static control, due to high variability in the static condition (**Figure 6A**). IL-6 (**Figure 6B**), CXCL8 (**Figure 6C**), and CXCL12 (**Figure 6D**) were also generally upregulated under 3.0 µm/s flow. Interestingly, the presence of naïve and activated CD4+ T cells did not generally impact the trends. CCL2 and IL-6 had similar trends with naïve CD4+ T cells compared to the control, but activated CD4+ T cells resulted in higher variability (**Figure 6A,B**). Notably, the presence of activated CD4+ T cells had the most significant impacts on CXCL8 (**Figure 6C**). Finally, CXCL12 secretion was variable in the presence of CD4+ T cells (**Figure 6D**).

**Figure 6.**
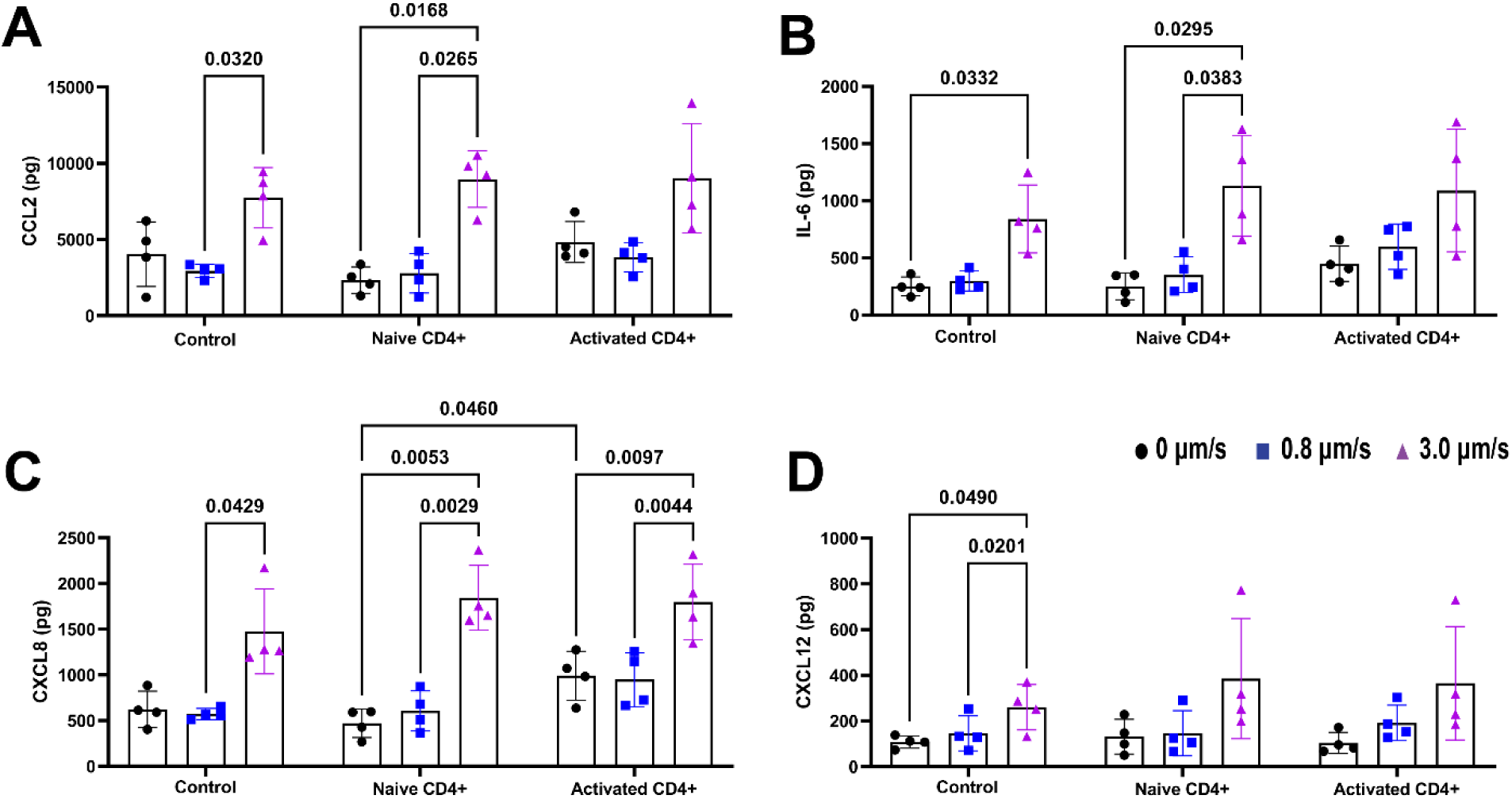
Inflammatory cytokine secretion is enhanced by 3.0 µm/s flow. Protein quantification of CCL2 (A), IL-6 (B), CXCL8 (C), and CXCL12 (D) was performed via Luminex in LN stroma models with and without CD4+ T cells. Each data point represents a biological replicate (n=3). Significance was determined by two-way ANOVA followed by Tukey’s t-test, with significant p values (<0.05) reported on each graph.

Finally, we degraded the LN stroma model hydrogel and examined the secretion of these chemokines within the matrix (**Supplemental Figure 10**). Interestingly, there were no significant differences in chemokine secretion in the control under flow, though it did trend downward. However, the presence of CD4+ T cells did result in some significant flow responses (**Supplemental Figure 10**).

Among these chemokines, CCL2 is an inflammatory marker known to be expressed by lymph node stromal cells in the T cell zone, with FRCs producing higher levels than LECs (58,59). In vitro, addition of CCL2 caused concentration-dependent enhanced chemotaxis (60,61). However, contrasting studies demonstrate impaired response to chemokines and homing to lymph nodes at low doses of CCL2 (62), but increased CD4+ and CD8+ T cell accumulation in LNs with higher CCL2 expression (63). IL-6 is known to enhance T cell motility (64), IFN-γ production (65), and cause decreased egress from the lymph node (66). The roles of CXCL8 and CXCL12 are less clear, but both are known to enhance T cell migration under certain conditions (67–71). However, the lack of increased secretion at 0.8 µm/s flow suggests that chemokines alone do not determine T cell egress in this LN stroma model.

### Flow magnitude-dependent responses in the lymph node stroma model

Ultimately, in this work we have built a model of the lymph node stroma and used it to test the response of the stroma, T cells, and their interactions to interstitial fluid flow. We demonstrated that varying interstitial fluid flow can drastically alter both morphology and protein secretion in the model. At 0.8 µm/s flow, changes to the LN stroma were less pronounced than at 3.0 µm/s, but still led to differences in T cell egress. Therefore, this flow rate may be utilized to answer questions about T cell migration in response to flow without the confounding factors of altered chemokines in the microenvironment. In contrast, we present the 3.0 µm/s flow rate as a model of inflammation in the lymph node stroma without any exogenous factors. This flow rate induced changes to FRC remodeling, LEC disruption, T cell behavior, and inflammatory chemokines (**Table 1**), and is ideal for answering questions around crosstalk in the lymph node during inflammatory states.

**Table 1.** All cell types in the LN stroma model are responsive to interstitial fluid flow. Changes to metrics relative to the control are denoted by ↑ for increase, ↓ for decrease, and – for unchanged.

| | Metric | + 0.8 $\mu\text{m/s}$ flow | + 3.0 $\mu\text{m/s}$ flow |
| --- | --- | --- | --- |
| LN Stroma Model | *FRC proliferation* | – | ↑ |
| | *FRC coverage* | – | ↑ |
|  | *LEC proliferation* | – | – |
| | *LEC disrupted junctions* | – | ↑ |
| | *LEC coverage* | – | ↓ |
|  | *Matrix remodeling genes* | – | – |
| | *IL-6 secretion* | – | ↑ |
| | *CCL2 secretion* | – | ↑ |
| | *CXCL8 secretion* | – | ↑ |
| | *CXCL12 secretion* | – | ↑ |
| + T cells | *FRC coverage* | – | – |
|  | *LEC disrupted junctions* | – | – |
| | *LEC coverage* | ↓ | ↓ |
| | *T cell egress* | ↓ | ↓ |
|  | *T cells in LN stroma* | – | – |
| | *IFNγ secretion* | – | ↑ |
| | *IL-6 secretion* | – | ↑ |
| | *CCL2 secretion* | – | ↑ |
| | *CXCL8 secretion* | – | ↑ |
|  | *CXCL12 secretion* | – | – |

## Conclusion

We have successfully established a spatially organized lymph node stroma model that is easily modified to test experimental questions regarding interstitial fluid flow and T cell retention. This work has established the complex effects of interstitial fluid flow within engineered lymph node stroma. Furthermore, it may suggest that other models of the immune system should also take fluid flow and other physical cues into account. The LN stroma model is designed with adaptability in mind, such as adding more immune cell types to increase complexity in future studies. Future work will aim to identify the mechanism behind flow-mediated changes to stroma and T cells and how crosstalk between the two under flow may alter response to antigens.

## Supplementary Material

Methodology and data for magnetic resonance imaging flow analysis and volumetric flow rate analysis is available in the supplementary material (Supplemental Figure 1 and 2). Additionally, assay information for gene expression targets is available in Table S1 and targets not shown in the main text can be found in Supplemental Figure 3. Representative images from second harmonic generation of the LN stroma models are found in Supplemental Figure 4. We examined how T cells would respond to interstitial fluid flow without the LN stroma, found in Supplemental Figure 5. Additional representative images and quantification from the LN stroma model in the presence of T cells can be found in Supplemental Figure 6-9. Finally, chemokine secretion into the hydrogel of the LN stroma rather than the media was quantified and displayed in Supplemental Figure 10.

## Acknowledgements

This work was supported by NIH-NIBIB NCATS grant U01 EB029127. JH was supported by the Virginia Tech Institute for Critical Technology and Applied Science (ICTAS). We would like to acknowledge Aileen C Suarez for their help with second harmonic generation imaging.

## Author Declarations

## Conflict of interest

The authors have no conflicts to disclose.

## Ethics Approval

FFPE tissue samples (AMSBIO) and LRS cones (STEMCELLTechnologies) containing cells were obtained de-identified with only subject sex and age provided and did not require ethics approval.

## Notes

### Competing Interest Statement

The authors have declared no competing interest.

