## Supplemental Materials for "Engineered human lymph node stroma model for examining interstitial fluid flow and T cell egress"

**Supplemental Methods:**

### MRI flow analysis

Tissue culture inserts with tissue engineered systems are secured in a 50 mL conical tube cap (**Figure S1A-C**). To allow interstitial flow, 100 μL of PBS is added beneath the tissue culture insert membrane. Tissue culture inserts are then stabilized and positioned in the MRI scanner. PE50 tubing was secured on the top of the tissue culture insert to infuse Gadolinium-DTPA via a syringe pump (Harvard Apparatus, Pump 11 Elite) later. Pre-contrast T1 images were taken as a baseline using a FLASH sequence with the following parameters: TE 3.0 ms, TR 150 ms, flip angle 30°, FOV 19.2 × 22.0 mm, matrix 192×220, slice number 9, slice thickness 0.8 mm, number of average 7, total scan time 3 mins. Then, 1:100 Gadolinium-DTPA (BioPal) was injected above the hydrogel at 1.0 ml/min flow rate. After 30 seconds, T1-weighted images were taken using the same parameters as baseline T1 images for 4 total post-contrast images. This dynamic series of contrast enhanced MRI images (DCE-MRI) captures the movement of the Gd-based contrast agent through the hydrogel over time. Pixel brightness corresponds to contrast agent concentration, allowing analysis of fluid movement within the substrate. We employ a physics-based method by solving the inverse problem of the diffusion-advection equation to estimate advection and diffusion parameters from observed pixel intensity changes, resulting in a vector fluid flow field of the hydrogel. To do this, we used “Lymph4D,” a previously published and openly available tool (1). This approach enables analysis of fluid flow properties such as divergence, as given by $div\ F = \nabla \cdot F = \partial F_x/\partial x + \partial F_y/\partial y$ where $F$ is the vector field within the gel. A lower divergence indicates more uniform flow and less spreading or expansion of the measured fluid.

### Volumetric flow rate analysis

PhotoHA-collagen hydrogels were prepared and pressure heads were added as described in the main text. After 2 hours, volume underneath the tissue culture insert is measured via pipettor. The liquid is replaced and the measurement is repeated at 4 hours and 6 hours. The volume is then used to calculate volumetric flow rate using the cross sectional area of the scaffold.

***Table S1. Gene expression targets and assay IDs.***

| Target | TaqMan Assay ID |
|---|---|
| CCL2 | Hs00234140_m1 |
| CXCL8 | Hs00174103_m1 |
| TGFbeta | Hs00998133_m1 |
| COL1A1 | Hs00164004_m1 |
| FN1 | Hs01549976_m1 |
| MMP2 | Hs01548727_m1 |

### Supplemental Figures:

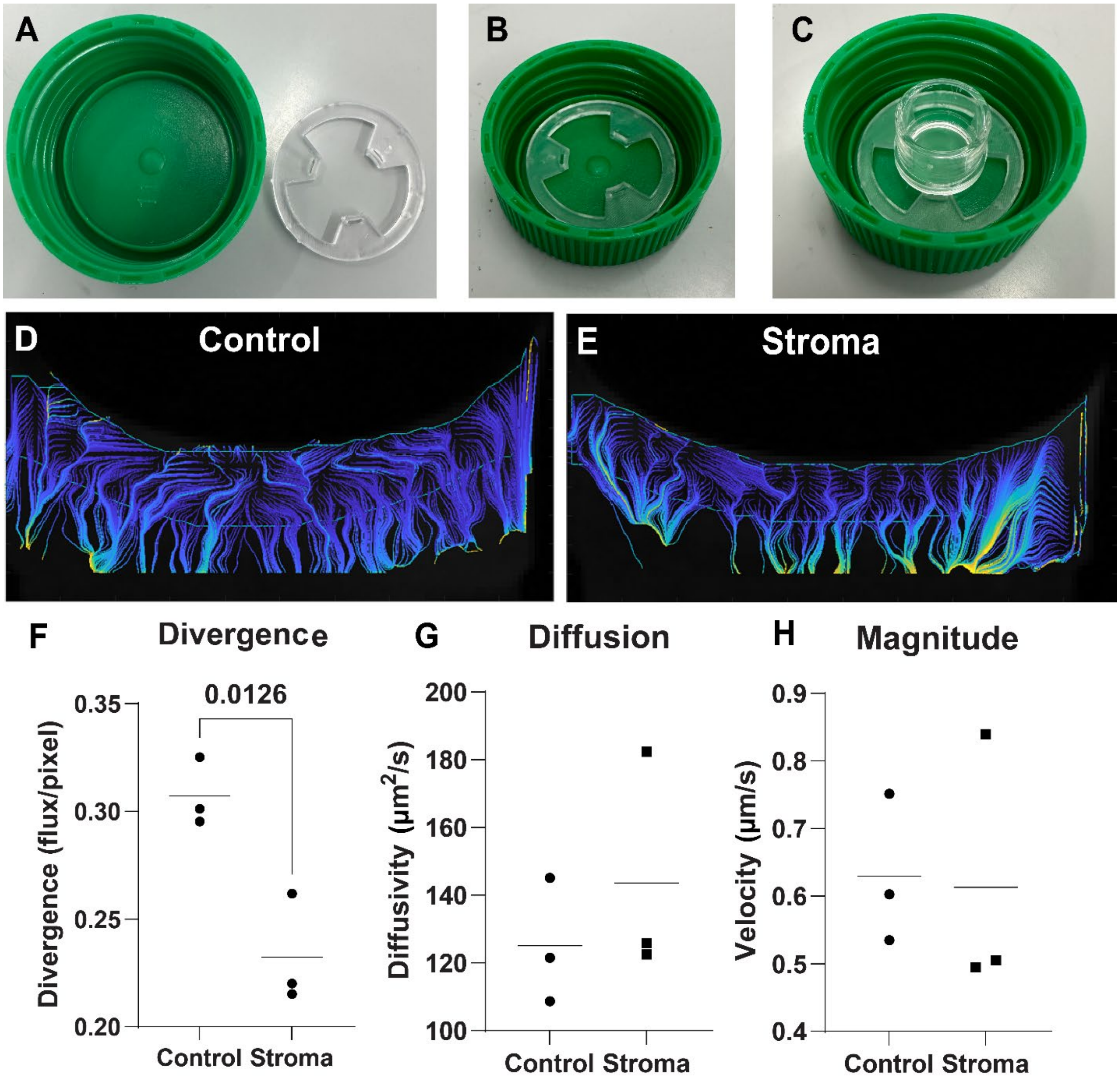


**Supplemental Figure 1. Magnetic resonance imaging for transport metrics of the LN stroma model.** A 3D printed insert with slots for tissue culture insert legs is created to the specifications of a 50 mL conical tube lid (A-C). The presence of LN stroma alters fluid transport in the hydrogel (D,E). Divergence of fluid is significantly decreased in the presence of LN stroma (F). Diffusion (G) and magnitude (H) are unchanged. Each data point represents a biological replicate (n=3). Significance was determined by students' t-test, with significant p values (<0.05) reported on the graph.

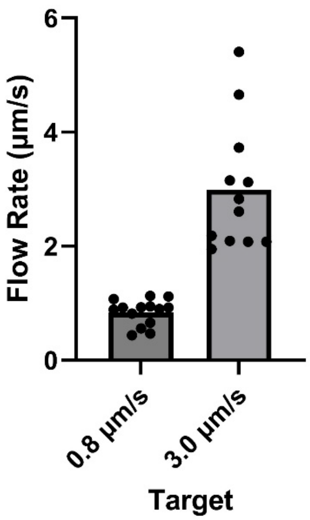


**Supplemental Figure 2.** Flow rate validation. Flow rates are quantified via volumetric measurement of media that passed through.

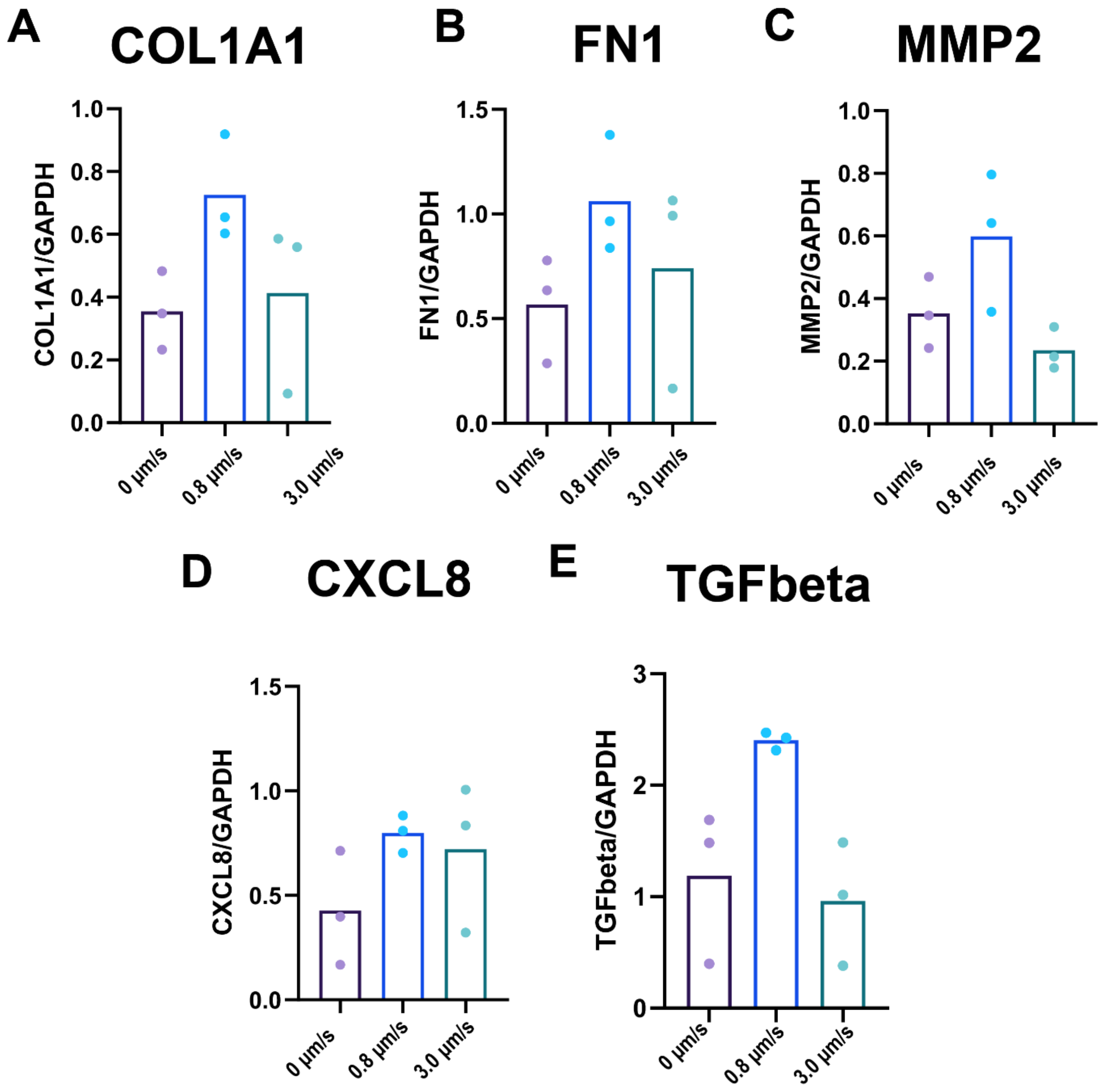


**Supplemental Figure 3.** Gene expression was analyzed in LN stroma models without T cells. For matrix remodeling, COL1A1 (A), FN1 (B), and MMP2 (C) are reported. For cytokines, CXCL8 (D) and TGFbeta (E) are reported. Each data point represents a biological replicate (n=3). Significance was determined by one-way ANOVA followed by Tukey's t-test, with significant p values (<0.05) reported on each graph.

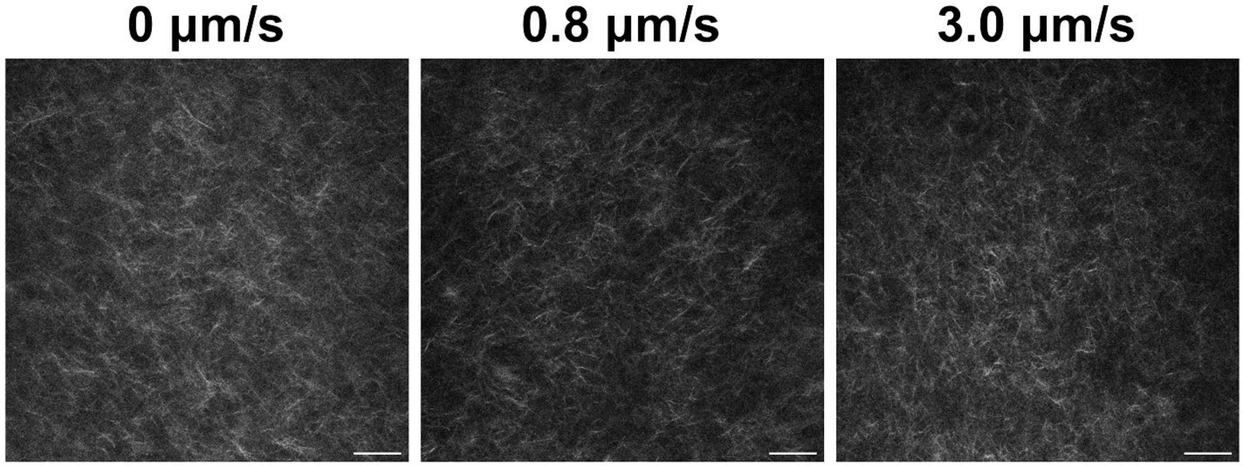


**Supplemental Figure 4.** Second harmonic generation imaging of collagen in the LN stroma models. Scale bar is 100 μm.

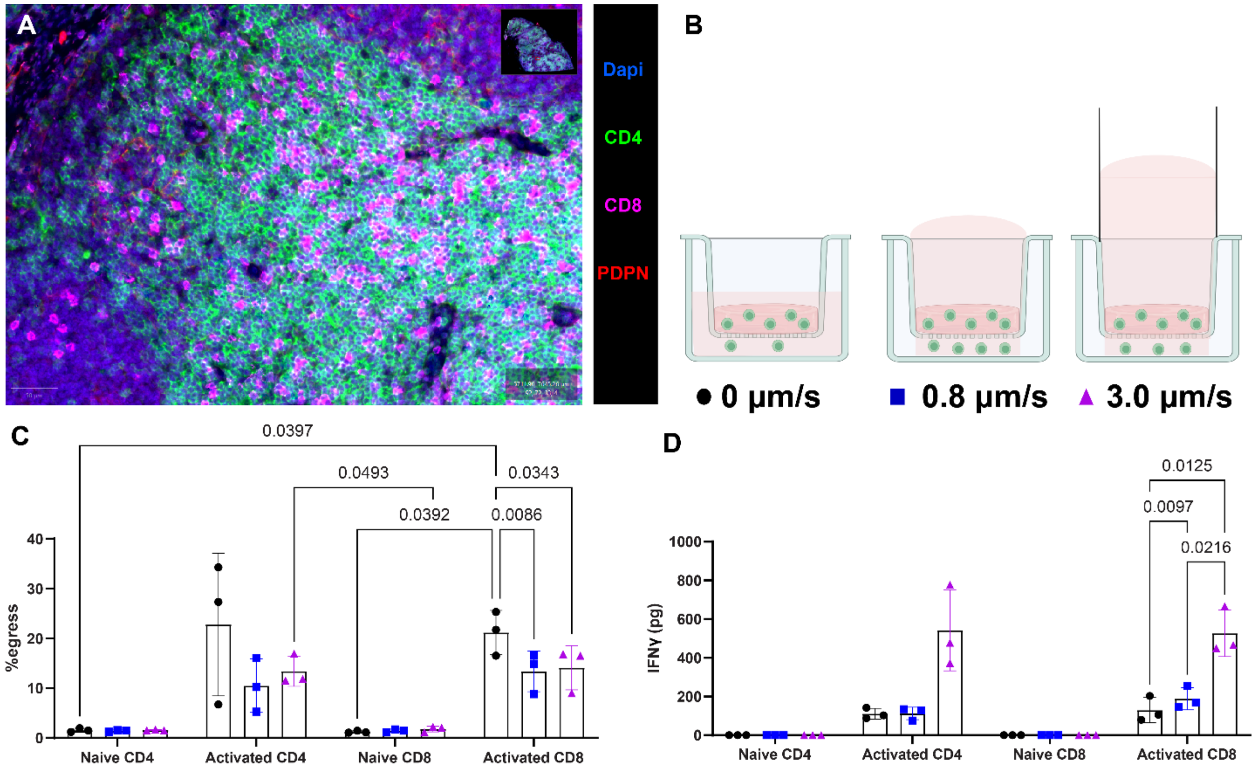


**Supplemental Figure 5. Activated T cell egress is decreased with interstitial fluid flow.** A human lymph node section demonstrates high density of CD4 and CD8 T cells (A). Scale bar is 50 μm. Schematic of T cells under varying pressure heads for IFF (B). Percent egress of naïve and activated CD4 and CD8 T cells is quantified and reported (C). Interferon-gamma secretion as determined by ELISA is quantified and reported (D). Each data point represents a biological replicate (n=3). Significance was determined by two-way ANOVA followed by Tukey's t-test, with significant p values (<0.05) reported on each graph.

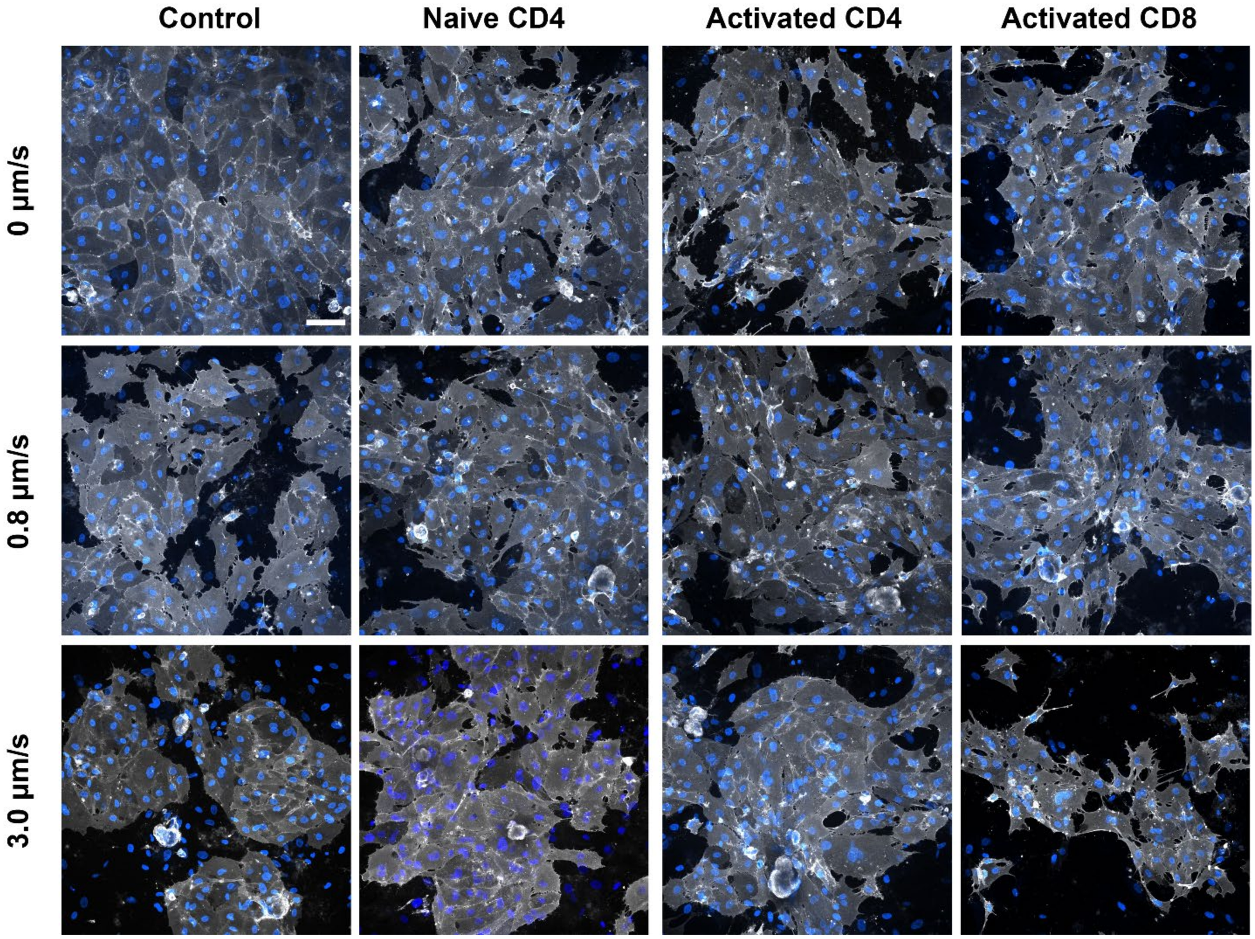


**Supplemental Figure 6.** Representative images of LEC monolayers in the presence of T cells. LECs are stained for CD31 (gray). Scale bars are 100 µm.

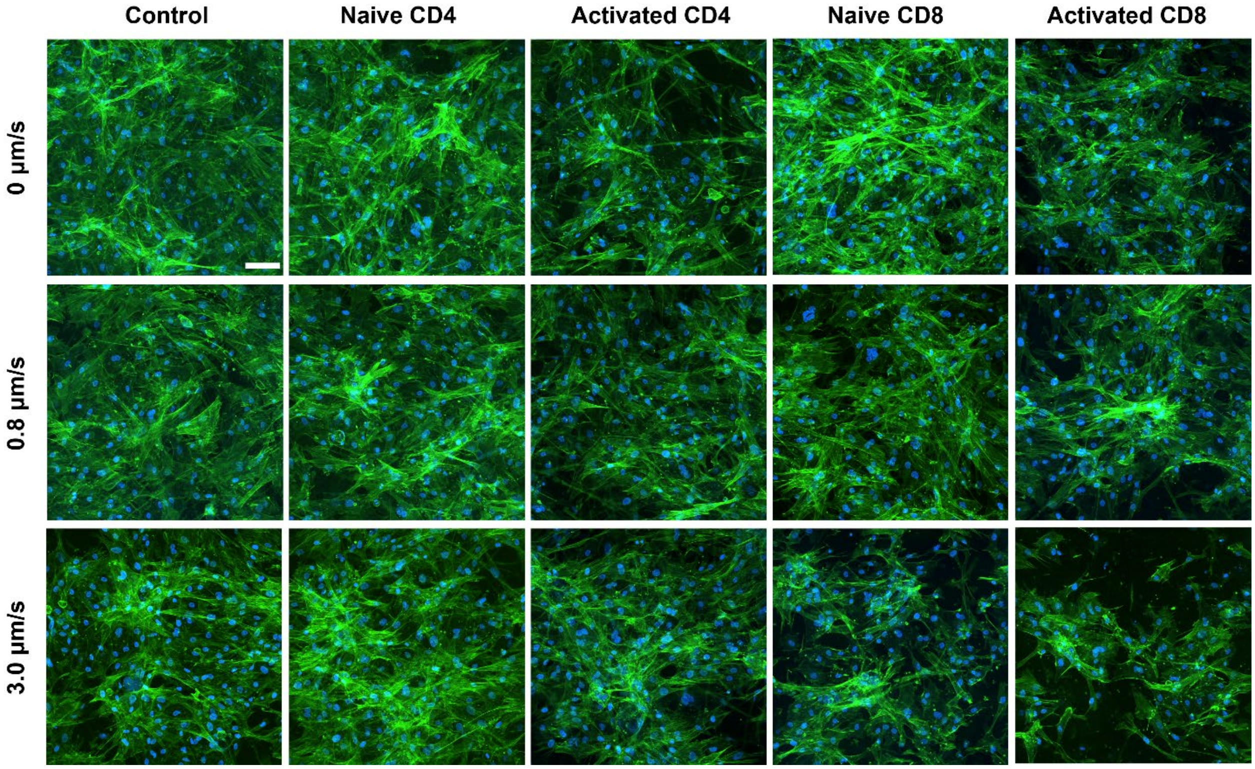


**Supplemental Figure 7.** F-actin staining on LEC monolayers. Representative images of LEC monolayers demonstrated fibroblast invasion. Scale bar is 100 µm.

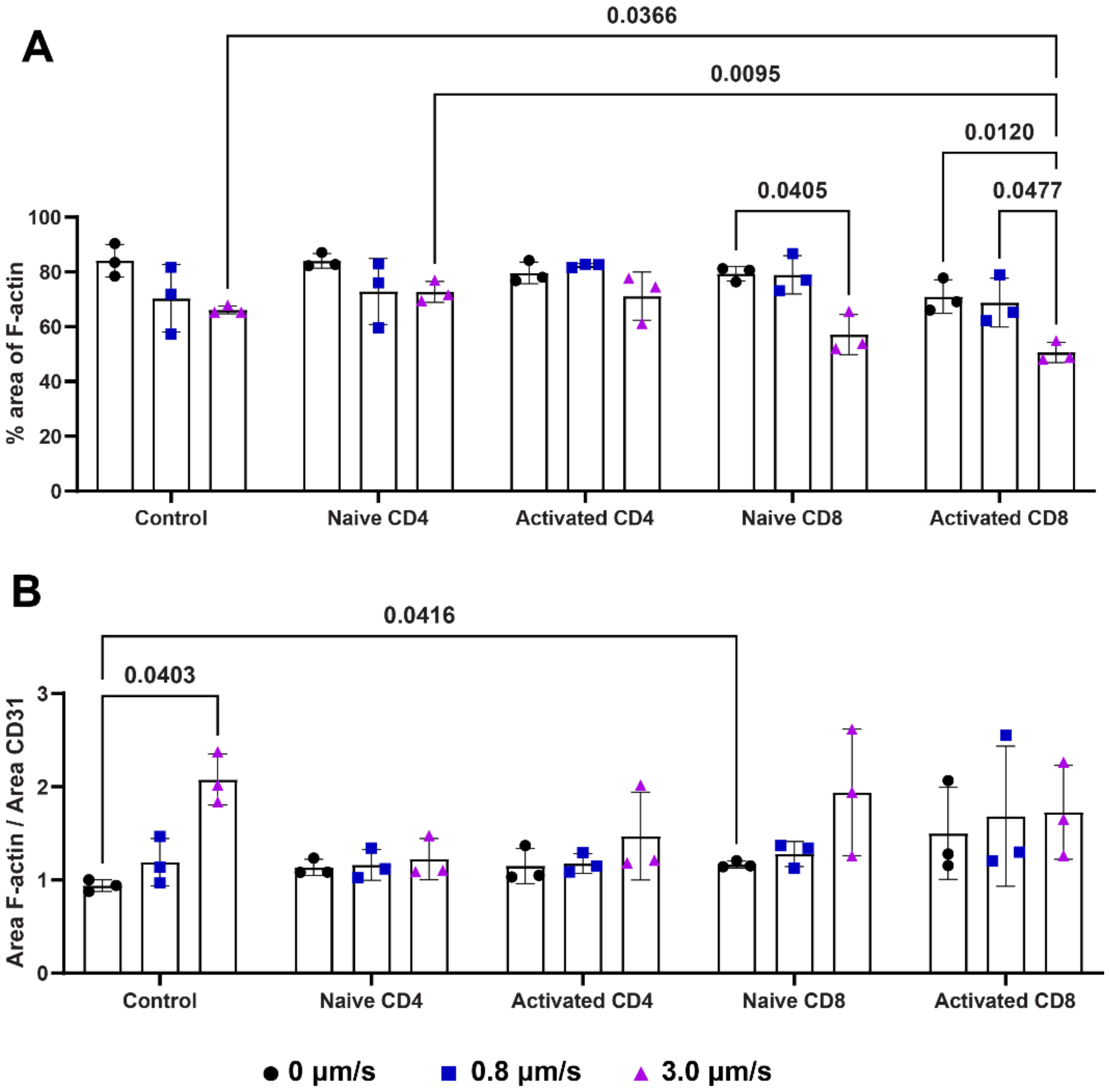


**Supplemental Figure 8.** The area of F-actin (green) on the monolayer (A) and the ratio of F-actin to CD31 (B) are reported. Each data point represents a biological replicate (n=3). Significance was determined by two-way ANOVA followed by Tukey's t-test, with significant p values (<0.05) reported on each graph.

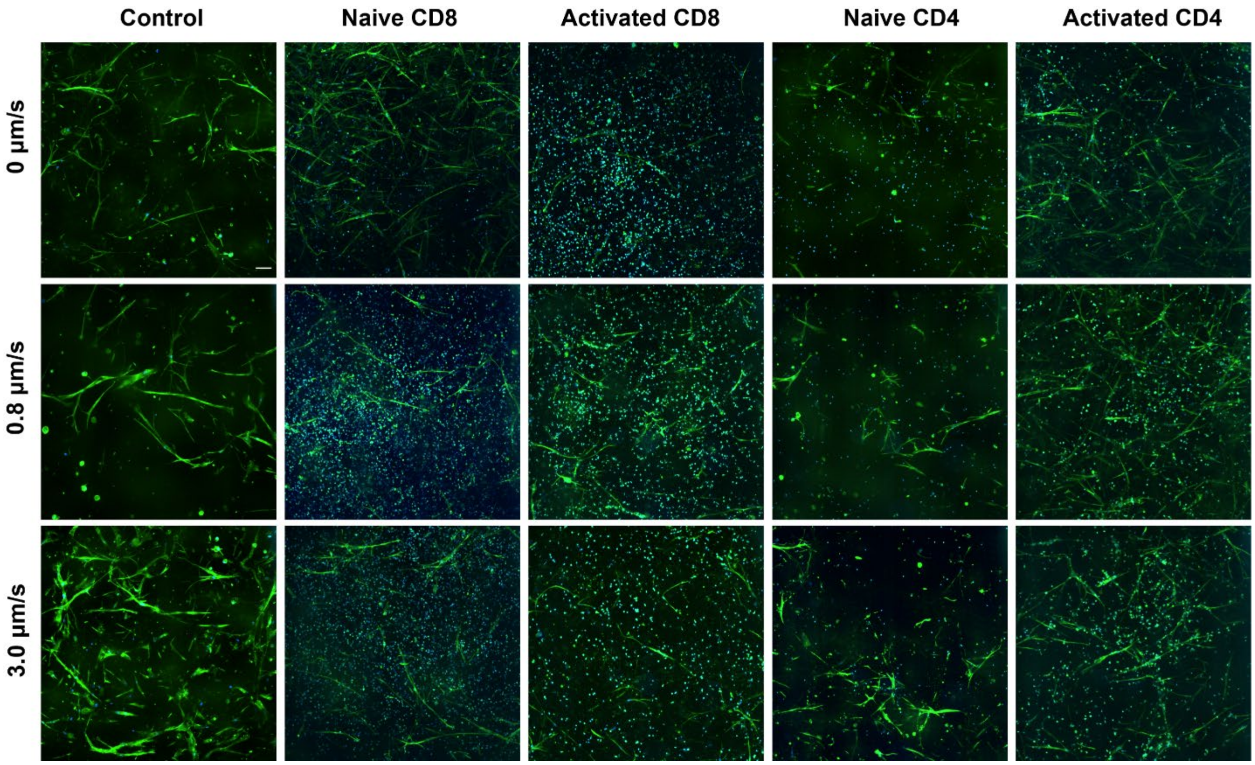


**Supplemental Figure 9.** Representative images of FRC networks (green) in the presence of T cells (blue). Scale bar is 100 µm.

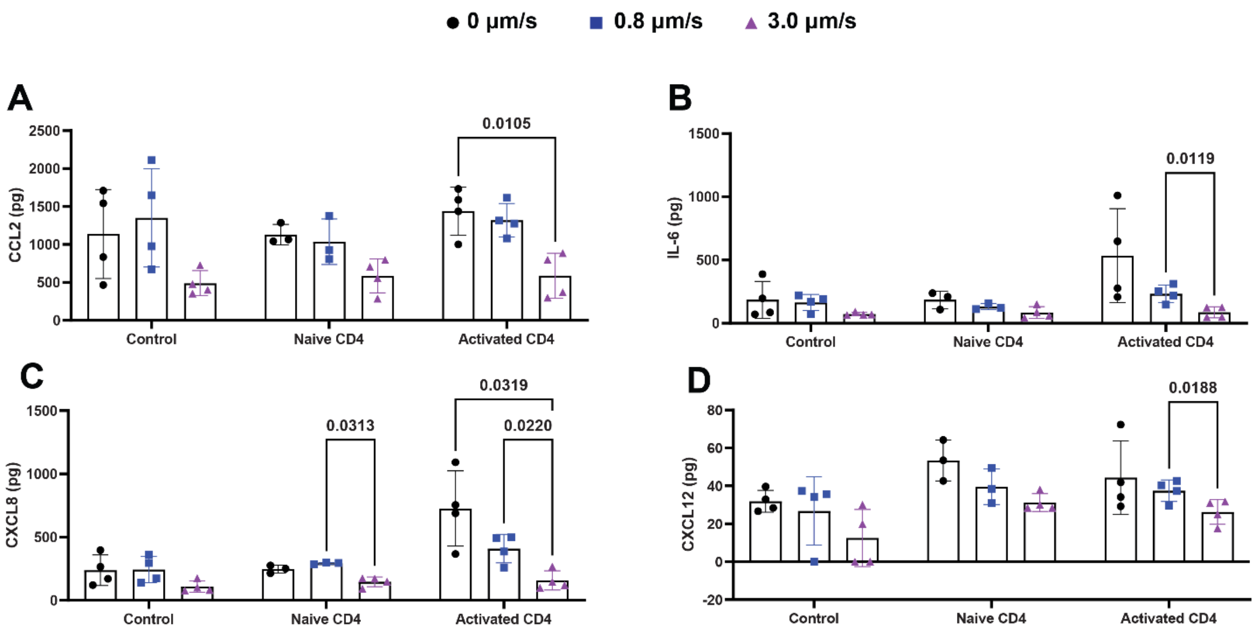


**Supplemental Figure 10. Chemokine secretion in the hydrogel.** Protein quantification of CCL2 (A), IL-6 (B), CXCL8 (C), and CXCL12 (D) was performed via Luminex in degraded LN stroma models with and without CD4+ T cells. Each data point represents a biological replicate (n=3). Significance was determined by one-way ANOVA followed by Tukey's t-test, with significant p values (<0.05) reported on each graph.
